# Immunogenicity of novel mRNA COVID-19 vaccine MRT5500 in mice and non-human primates

**DOI:** 10.1101/2020.10.14.337535

**Authors:** Kirill V. Kalnin, Timothy Plitnik, Michael Kishko, Jinrong Zhang, Donghui Zhang, Adrien Beauvais, Natalie G. Anosova, Timothy Tibbitts, Joshua M. DiNapoli, Po-Wei D. Huang, James Huleatt, Deanne Vincent, Katherine Fries, Shrirang Karve, Rebecca Goldman, Hardip Gopani, Anusha Dias, Khang Tran, Minnie Zacharia, Xiaobo Gu, Lianne Boeglin, Sudha Chivukula, Ron Swearingen, Victoria Landolfi, Tong-Ming Fu, Frank DeRosa, Danilo Casimiro

**Affiliations:** Sanofi Pasteur, 38 Sidney Street, Cambridge, MA 02139; Sanofi Pasteur, Discovery Dr., Swiftwater, PA 18370; Translate Bio, 29 Hartwell Ave, Lexington, MA 02421; Sanofi Pasteur, 1541 AV Marcel Mérieux, 69280 Marcy l’Etoile, France; Yoh Services LLC, 38 Sidney Street, Cambridge, MA 02139

**Keywords:** COVID-19, SARS-CoV-2, vaccine, mRNA, LNP, BALB/c mice, cynomolgus macaques, immunogenicity, neutralization potency, neutralization, microneutralization, ELISA

## Abstract

An effective vaccine to address the global pandemic of coronavirus disease 2019 (COVID-19) is an urgent public health priority^1^. Novel synthetic mRNA and vector-based vaccine technologies offer an expeditious development path alternative to traditional vaccine approaches. Here we describe the efforts to utilize an mRNA platform for rational design and evaluations of mRNA vaccine candidates based on Spike (S) glycoprotein of Severe Acute Respiratory Syndrome Coronavirus 2 (SARS-CoV-2), the virus causing COVID-19. Several mRNA constructs expressing various structural conformations of S-protein, including wild type (WT), a pre-fusion stabilized mutant (2P), a furin cleavage-site mutant (GSAS) and a double mutant form (2P/GSAS), were tested in a preclinical animal model for their capacity to elicit neutralizing antibodies (nAbs). The lead 2P/GSAS candidate was further assessed in dose-ranging studies in mice and *Cynomolgus* macaques. The selected 2P/GSAS vaccine formulation, now designated MRT5500, elicited potent nAbs as measured in two types of neutralization assays. In addition, MRT5500 elicited T_H_1-biased responses in both mouse and non-human primate species, a result that helps to address a hypothetical concern regarding potential vaccine-associated enhanced respiratory diseases associated with T_H_2-biased responses. These data position MRT5500 as a viable vaccine candidate for clinical development against COVID-19.

## Introduction

SARS-CoV-2, previously known as the 2019-novel coronavirus (2019-nCoV)^2^, is a β-coronavirus with a yet-to-be defined zoonotic origin. The first cases of human infection with severe acute respiratory syndrome (SARS) were reported in December 2019 in China^3^, and later named coronavirus disease 2019 (COVID-19)^4^. In contrast to SARS-CoV-1 virus which caused an outbreak in 2002, SARS-CoV-2 has gained high capacity for human-to-human transmission and quickly spread worldwide. It has caused over 34 million cases of confirmed infection and more than 1,000,000 deaths in 188 countries (https://www.worldometers.info/coronavirus). An effective vaccine is urgently needed to address this global pandemic.

Coronavirus is an enveloped RNA virus, and the viral spike (S) protein on the virion envelope is essential for infection and is the target for host antiviral antibodies^5,6^. The receptor for SARS-CoV-2 is angiotensin-converting enzyme 2 (ACE2), a metalloprotease that also serves as the receptor for SARS-CoV-1^7^. Most of the COVID-19 vaccine candidates reported are focused on a pre-fusion-stabilized S protein, either as recombinant protein with adjuvant or delivered from viral vectors or as DNA or mRNA vaccines ^8–15^. The pre-fusion-stabilized version of SARS-CoV-2 S-protein contains two proline substitutions (2P), at amino acid positions 986 and 987, located near the apex of the central helix and heptad repeat 1^16^. Structural studies reveal that the pre-fusion stabilized S closely resembles native S protein on the virion surface; a structure targeted by many reported effective neutralizing antibodies^17–19^. Moreover, the vaccine premises are based on the prior work of MERS-CoV, SARS-CoV and HCoV-HKU1 S proteins presented in pre-fusion conformations^20–22^. The ability of S-2P-based vaccines to elicit neutralizing antibodies has been demonstrated ^8–10 23,24^.

There is a unique feature of SARS-CoV-2 S protein which possesses a polybasic furin cleavage site at the junction of S1 and S2 subunits. This feature is believed to have emerged during viral transmission from zoonotic host to human^25–27^, and is key to SARS-CoV-2 high transmissibility in humans^28,29^. Although robust SARS-CoV-2 infection of human lungs requires a multibasic cleavage site^30^, interestingly, both cleaved and uncleaved versions of S protein co-exist on virions purified from viral culture on Vero cells^31,32^. Thus, it remains unclear how the cleavage provides an advantage for viral transmission. Also, from a vaccine design perspective, one may speculate that furin cleavage site may result in subtle conformational changes in the trimerized S protein, potentially favoring its interaction with ACE2^26^.

These unanswered questions led us to design various forms of S protein constructs involving both 2P and cleavage site, referred to herein as GSAS mutations. These GSAS constructs were first evaluated for immunogenicity in mice. The 2P/GSAS S mRNA encapsulated in a cationic lipid nanoparticle (LNP) formulation, designated as MRT5500, was subsequently selected for further evaluation. Here we report the results of preclinical evaluation of MRT5500 in mice and non-human primates (NHPs). MRT5500 was administered twice via the intramuscular route (IM) at a three-week interval in both animal models. Results demonstrated that MRT5500 elicited potent neutralizing activity and a T_H_1-biased response in both species. The ability of this vaccine to induce both humoral and cell-mediated antiviral responses identifies MRT5500 as a promising clinical vaccine candidate.

## Results

### Design and selection of mRNA constructs

SARS-CoV-2 S protein, a 1273 amino acid glycoprotein, is expressed and stabilized as a membrane anchored homo-trimer^6^. The receptor binding domain (RBD) has been identified as the critical component to initiate virus attachment to ACE2, a cellular receptor for viral infection^33^. Interestingly, the RBD is present in both up and down configurations in the pre-fusion form of S protein, and the up position has been speculated as the prerequisite for interaction with ACE2^6,31^. The furin cleavage at the S1/S2 boundary of SARS-CoV-2 S occurs during viral biosynthesis^34^. It is postulated that transition and adaptation to the human host resulted in the acquisition of a furin protease site in the S protein of SARS-CoV-2, which is a unique feature discriminating this virus from SARS-CoV-1 and other SARS-related-CoVs^26^. Approximately 45% of the total S protein monomers presented within intact SARS-CoV-2 virions have been reported as cleaved at the furin cleavage site^31^; however, it is not clear which form is favored by the virus to facilitate the fusion process^26,34–36^.

The COVID-19 vaccine hypothesis has been centered around induction of neutralizing antibodies (nAbs) that either block the interaction of the RBD with ACE2, or that prevent the fusion process involving S protein transition from pre- to post-fusion conformation^37,38^. Although the pre-fusion conformation is known to be critical for eliciting a neutralizing response^18,19^, the impact of the furin cleavage site in eliciting neutralizing antibodies requires additional studies. To test the potential contribution of this site, we mutated the furin cleavage site, composed of the polybasic residues RRAR, to GSAS from amino acid position 682 to 685^30,35^. Four constructs were synthesized as mRNA to represent either wild type (WT), stabilized pre-fusion mutant (2P) ^20^, furin cleavage site mutant (GSAS) or a double mutant (2P/GSAS) of SARS-CoV-2 S gene. These constructs were transfected into a human cell line and their expression levels were verified by Western Blot (**Fig. 1a**). As expected, endogenous cleavage of WT and 2P constructs, but not GSAS or 2P/GSAS proteins, was observed (**Fig. 1a**), which yielded a band of approximately 90 kDa representing S2.

**Figure 1:**
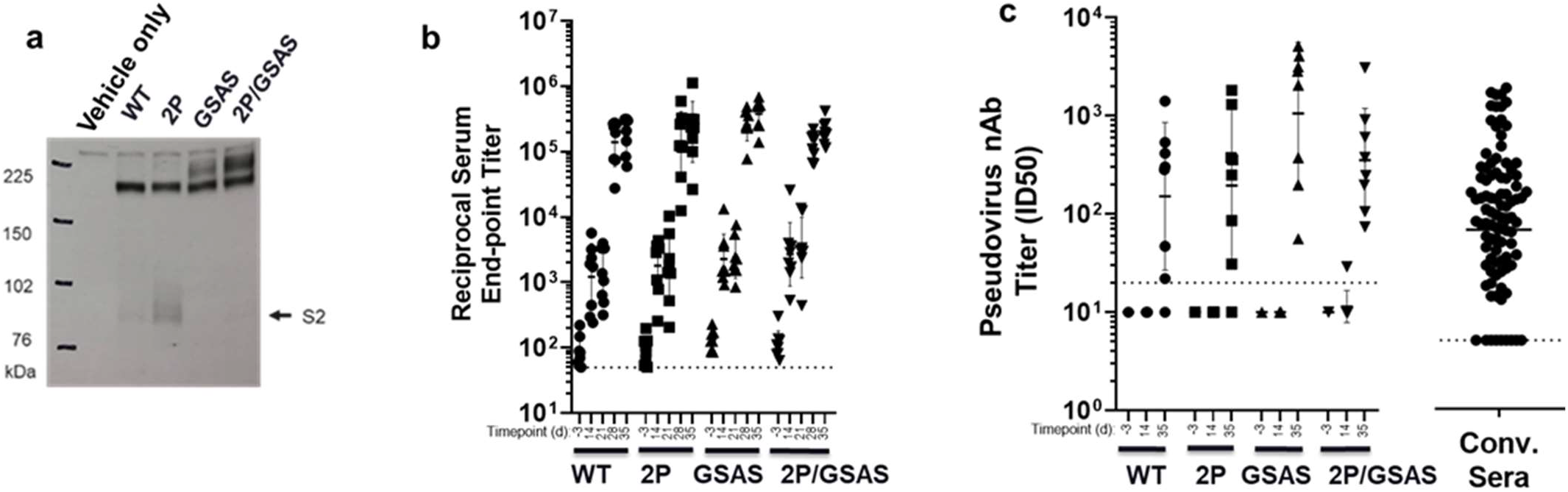
Comparison of S antigen constructs. (**a**) In vitro expression of S SARS-CoV-2 protein was assessed in Western blot analysis HEK293 cells were transfected with 1 μg mRNA construct of control diluent (Mock), wild type (WT), stabilized pre-fusion mutant (2P), cleavage site mutant (GSAS) or double mutant containing both mutations (2P/GSAS) and the respective expressed S proteins were detected with rabbit anti-SARS-CoV spike protein polyclonal antibody (NB100-56578, Novus Biologicals). (**b**) Serum antibodies reactive to S antigen and (**c**) serum neutralizing titers in mice immunized with mRNA vaccines. BALB/c female mice (n=8) were immunized at D0 and D21 with 0.4 μg of WT, 2P, GSA, 2P/GSAS mRNA vaccine formulations. Sera samples at indicated timepoints were tested for reactivity to recombinant S protein in ELISA or tested in a pseudovirus neutralization assay. The 50 % inhibitory dilution titers (ID_50_) were calculated as the reciprocal of the dilution that reduced the number of virus plaques in the test by 50%. Each dot represents an individual serum sample and the line represents the geometric mean with standard deviation for the group. the dotted line below for each panel represents the lower limit of assay readout.

In order to determine the potential impact of 2P and GSAS mutations on immunogenicity, we formulated each of the four mRNA constructs within a lipid nanoparticle (LNP), which has been designed for efficient delivery of mRNA vaccines^39^. BALB/c mice were administered two immunizations at a 0.4 μg dose of each of four formulations at a three-week interval. Binding antibody activities in the serum samples were assessed via Enzyme-Linked Immunosorbent Assay (ELISA) (**Fig. 1b**). All four vaccines demonstrated similar levels of binding antibodies 14 days after the first vaccination, and the responses were further enhanced one week after the second dose at day (D) 28. On D35, the IgG geometric mean titers (GMTs) for WT, 2P, GSAS and 2P/GSAS groups were 184343, 200896, 379653 and 201080 respectively. There were no statistically significant differences among those GMT titers.

To understand the potential impact of these mutations on nAbs titers, we tested the ability of immune sera to neutralize the infectivity of GFP reporter pseudoviral particles (RVP) in HEK-293T cells stably over-expressing human ACE2 ^40^. RVPs expressed antigenically correct SARS CoV-2 S protein and GFP reporter genes on lentiviral (HIV) core and were capable of a single round of infection. Pseudoviral neutralization assay (PsVNa) allowed the determination of serum dilution which can achieve 50% inhibition of RVP entry (ID_50_; see Materials and Methods). Contrary to binding antibodies which could be detected at D14 after the first immunization, the neutralizing antibody response could only be detected after the second immunization. Also noted, the spread of the nAb titers within each group were more pronounced when compared to binding antibody titers, with 95% confidence intervals overlapping each other. On D35, the GMTs for pseudoviral (PsV) nAb titers were 152 for WT, 195 for 2P, 1005 for GSAS and 354 for 2P/GSAS. The neutralizing potential of the GSAS variant was trending slightly higher than 2P/GSAS.

Another important observation is that ELISA titers were not consistently predictive of neutralizing titers by PsVNa. Some mice in the WT and 2P groups did not seroconvert in the neutralization assay but their endpoint ELISA titers were comparable to the other animals in the group which demonstrated neutralizing activity. We therefore placed greater emphasis on PsVNa titer for continuing candidate evaluation. Considering the trend towards higher PsVNa observed for the GSAS constructs as well as the expected importance of the pre-fusion conformation, we selected the double mutant 2P/GSAS/LNP formulation, referred to as MRT5500, for further preclinical evaluations.

### Serological evaluations of MRT5500 in mice and NHPs

The selected MRT5500 formulation was evaluated in both mouse and NHP studies with a range of doses covering more than 10-fold titration. The hypothesis for this study is that S-specific antibodies blocking viral infection are key for protection, and our evaluation therefore focused on serological responses against SARS-CoV-2 S, with a particular emphasis on neutralizing titers post vaccination ^8,9,41^. Four dose levels in mice were assessed, ranging from 0.2 to 10 μg per dose. As expected, MRT5500 induced dose-dependent S-specific binding antibodies and neutralizing antibodies in mice (**Fig. 2**). PsVNa titers were detected in the higher dose groups (5 μg, 10 μg) after one vaccination, within the titers being more pronounced after the second vaccination at D21 (**Fig. 2b**). The PsVNa GMTs were 534, 5232, 9370 and 7472 at D35 for the 0.2, 1.0, 5.0 and 10.0 μg dose groups, respectively. There were no statistically significant differences in PsV neutralization titers on D35 between 1, 5 and 10 μg groups (**Suppl. Table 2**), suggesting a dose-saturation effect beyond 1 μg in mice. We also demonstrated that the peak PsV titers (D35) in mice were significantly different from the titers observed in a panel of 93 convalescent sera from COVID-19 patients (**Suppl. Fig. 4**).

**Figure 2:**
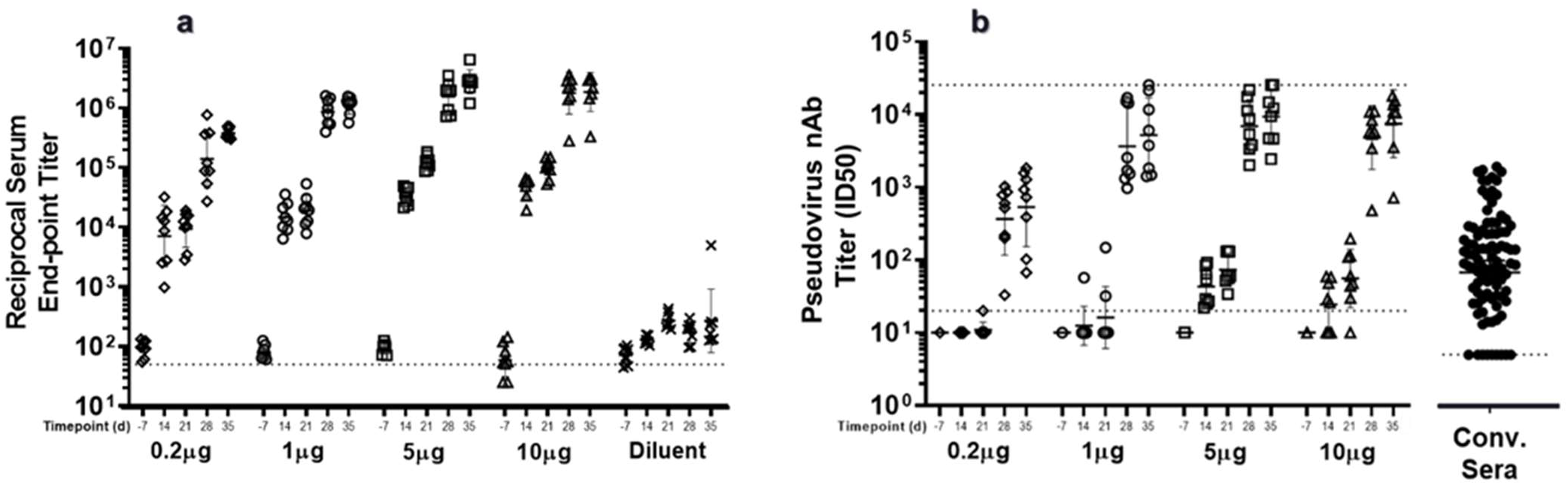
Serological evaluation of MRT5500 formulation in mice. Groups of BALB/c mice (n=8) were immunized at D0 and D21 with 0.2, 1.0, 5.0 or 10.0 μg dose of MRT5500 formulation. Serum samples at the indicated time were tested in ELISA (**a**) and PsVNa (**b**). Each symbol represents a serum sample and the line is the geometric mean with standard deviation of the group. The dotted line in each panel represents the lower limit of assay detection. PsV neutralization titers (nAb) of the human convalescent serum panel (n=93) were defined in separate experiment and shown in the same scale on Y-axis as other samples.

In NHPs, we evaluated three dose levels: 15, 45 or 135 μg per dose. After the first immunization, nearly all NHPs (10 out of 12) developed antibodies reactive to recombinant S protein in ELISA, and the titers were further enhanced after a second immunization at D35 (**Suppl. Fig.1**) with all NHPs demonstrating high titers of nAbs. The neutralization potency was assessed by two methods: PsVNa (**Fig. 2a**) and microneutralization (MN) assay (**Fig. 2b**). In both assays, a dose-dependent increase in neutralization titer was observed, with GMTs on D35 of 924 for 15 μg, 961 for 45 μg, and 2871 for 135 μg in PsVNa. The MN GMTs followed a similar trend, with titers of 555 for 15 μg, 719 for 45 μg and 1877 for the 135 μg group. Despite the observed trend towards higher titers with increasing dose, the differences between groups was not statistically significant for either MN or PsV neutralization titers.

Although we have used two assays to measure the neutralizing potency, the results from both assays were highly correlated (**Suppl. Fig. 3** and **Suppl. Table 1**). Regardless of the dose level tested, D35 PsV and MN titers were approximately 130-fold higher than those of pre-immune animals. Furthermore, the observed PsV and MN titers were significantly higher from titers observed in a panel of 93 convalescent sera from COVID-19 patients (**Suppl. Fig. 5**).

### T-cell profiles of the selected mRNA formulation in NHPs

Vaccine associated enhanced respiratory disease (VAERD) has been a safety concern for COVID-19 vaccines in development, although the concern at this stage is only a theoretical one^1^. This phenomenon has been reported for whole-inactivated virus vaccines against measles and respiratory syncytial virus (RSV), which were tested in the 1960s (cit by ^1^), and one of the disease hypotheses implicates the biased production of T_H_2 cytokines (IL-4, IL-5, IL-13) by antigen-specific CD4 T cells. A similar association between a T_H_2 profile and disease enhancement has been reported for an inactivated SARS-CoV-1 vaccine in mice ^42^. Furthermore, less severe cases of SARS were associated with accelerated induction of T_H_1 cell responses^43^, whereas T_H_2-biased responses have been associated with enhancement of lung disease following infection in mice parenterally vaccinated with inactivated SARS-CoV viral vaccines^42,44^. Similar phenomena have been observed in humans. For example, a SARS-CoV-2-specific cellular response was associated with severity of disease: recovered patients with mild COVID-19 illnesses demonstrated high levels of IFN-γ induced by SARS-Cov-2 antigens, while severe pneumonia patients showed significantly lower level of this cytokine^45^. Thus, it is important to understand the T cell profiles induced by MRT5500.

T cell cytokine responses were tested in NHPs three weeks after the second vaccination. Cytokines induced by restimulation with the pooled SARS CoV-2 S protein peptides were assesses in PBMCs on D42 by the IFN-γ (T_H_1 cytokine) and IL-13 (T_H_2 cytokine) ELISPOT assays. The majority of animals in three dose level groups tested (10 out of 12) demonstrated presence of IFN-γ secreting cells, ranging from two to over 100 spot-forming cells per million PBMCs. A dose-response was not observed as the animals in the lower and higher dose level groups showed comparable frequencies of IFN-γ secreting cells. In contrast, presence of IL-13 cytokine secreting cells was not detected in any of the groups tested and at any dose level, suggesting induction of a T_H_1-biased cellular responses (**Fig. 4**). These data presented clear evidence for lack of T_H_2 response to S antigen following vaccination in NHPs.

**Figure 3:**
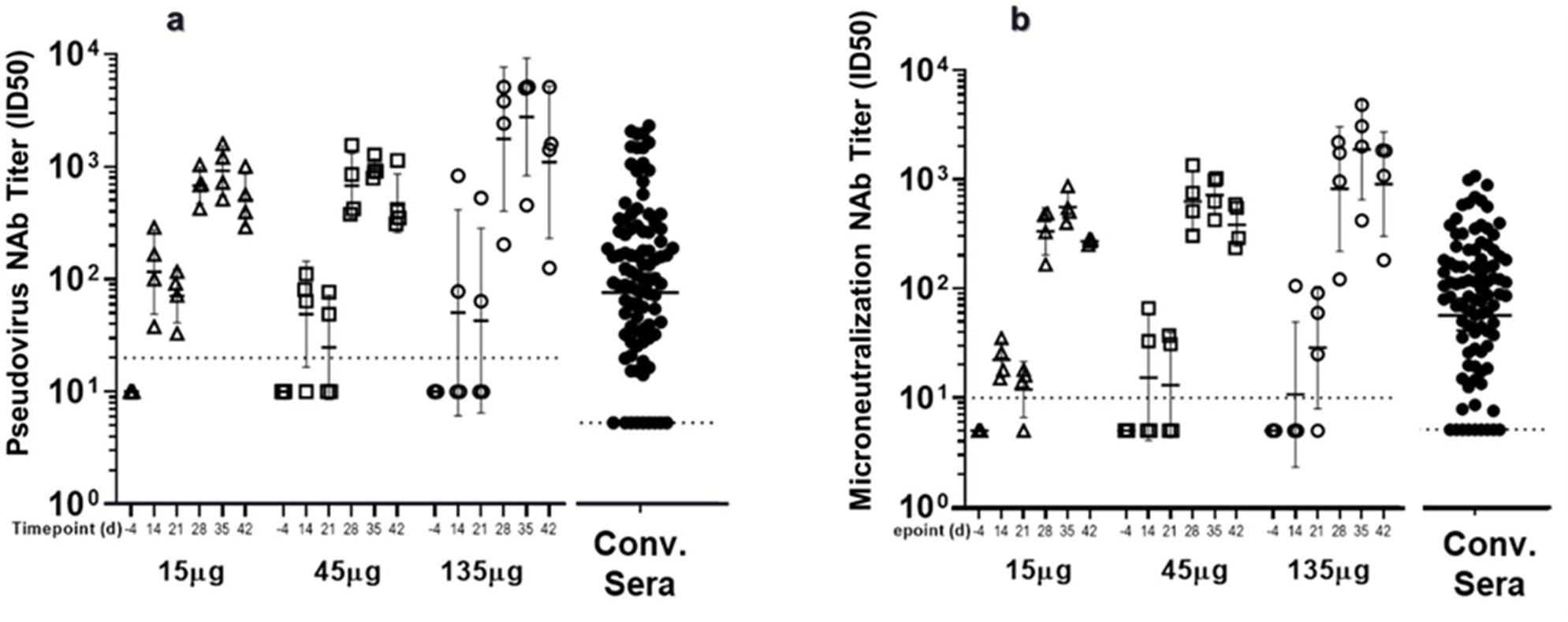
Neutralizing titers in NHPs vaccinated with MRT5500 formulation. Groups of cynomolgus macaques (n=4) were vaccinated with MRT5500 at 15, 45 or 135 μg per dose at D0 and D21, and serum samples collected at the indicated timepoints were tested in PsVNa (**a**) and MN assay (**b**). Each symbol represents an individual sample and the line geometric means for the group. The neutralization titer of the sample, shown as ID_50_, was defined as the reciprocal of the highest test serum dilution for which the virus infectivity was reduced by 50% when compared to the assay challenge virus dose. PsV and MN neutralization titers (NAb) of the human convalescent serum panel (n=93) were defined in separate experiment and shown in the same scale on Y-axis as other samples.

**Figure 4:**
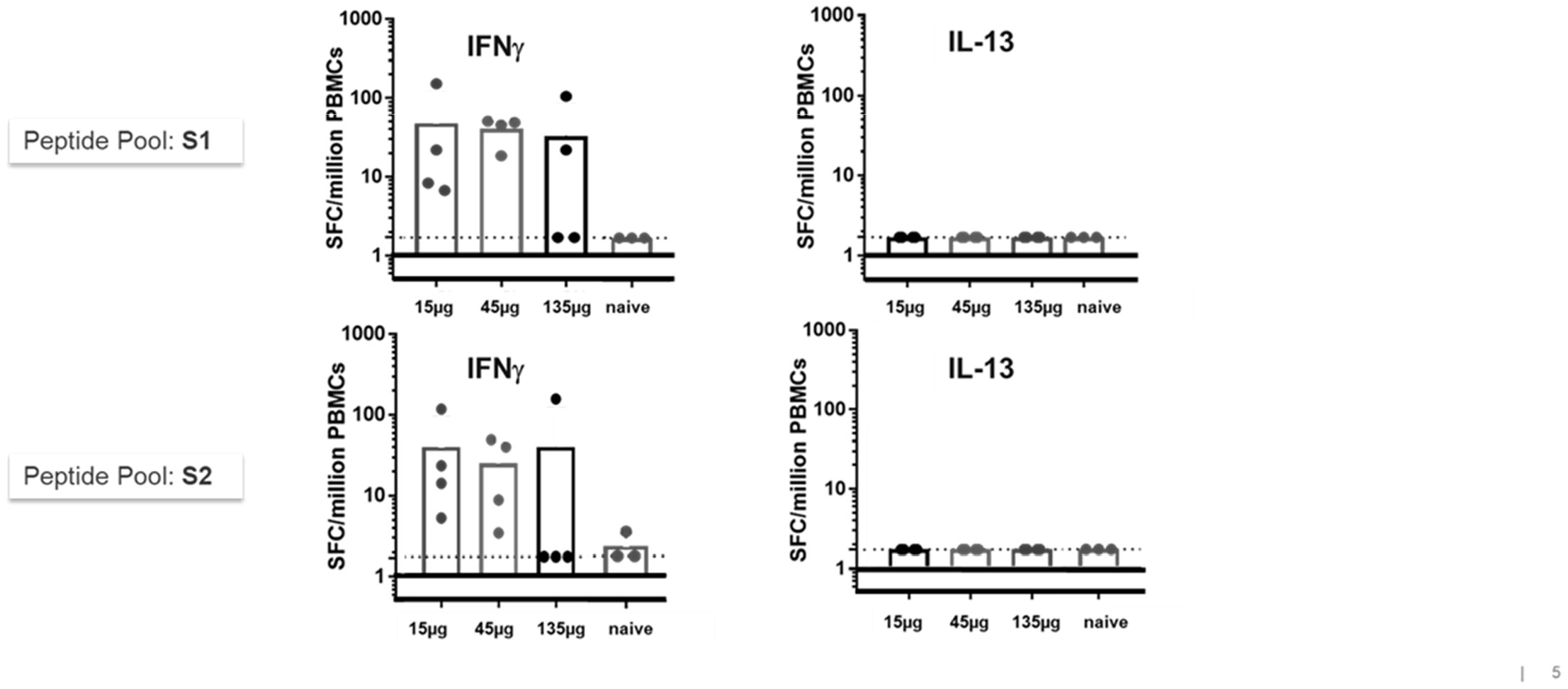
Assessment of T-cell responses in NHPs vaccinated with MRT5500. PBMCs collected at D42, 21 days post the second vaccination, were incubated overnight with the SARS-Cov-2 S-protein peptide pools representing the entire S open reading frame. The frequencies of PBMC secreting IFNγ (left panels) or IL-13 (right panels) were calculated as spots forming cells (SFC) per million PBMC. Each symbol represents an individual sample, and the bar represent the geometric mean for the group. The dotted line represents the lower limit for detection.

Similar assessment of cellular immune responses was performed in immune splenocytes in BALB/C mice on D35. ELISPOT was conducted in the 5 and 10 μg dose groups in Fig. 2. Although BALB/c mice have strong tendency for T_H_2 biased immune responses, following re-stimulation with the S protein peptide pools splenocytes from the MRT5500 immunized mice secreted predominantly IFNγ while IL-5 responses were marginal, suggesting considerable T_H_1 bias (**Supplementary Fig.2**). Thus, MRT5500 vaccination elicited predominantly T_H_1-biased responses in both animal species.

## Discussion

mRNA-based vaccine development provides a rapid pathway for effective evaluation of multiple vaccine construct designs which we employed for our initial evaluation of S antigen mRNA vaccine candidates against SARS-CoV-2. For any vaccine intended to generate antibody-mediated immunity, delivering a conformationally correct protein is critical^1^. Lessons have been learned from RSV F protein where the post-fusion form elicited poor neutralizing antibodies, albeit extremely immunogenic in humans^46^, and the post-fusion form F antigen vaccine has failed to provide any protection against RSV infection^47^. Thus, our focus in this study was to identify mutations that could stabilize the pre-fusion form of S antigen upon expression. In contrast to the other S antigen mRNA vaccines under evaluations^9,10,23,24,48^, we have incorporated a unique mutation at cleavage site GSAS, in addition to 2P, which has enhanced features to lock the S protein in the pre-fusion form. There were two considerations for this design. First, it is not known whether 2P mutations alone, located at the apex of the central helix and heptad repeat 1, are sufficient for locking the S antigen in the pre-fusion form. Second, it has been hypothesized that cleavage of S antigen into S1 and S2 subunits is part of the transition from the pre-fusion to post-fusion form during viral entry^19^. Thus, by blocking the furin cleavage site, we have added another layer for prevention of pre-fusion to post-fusion conversion.

The two GSAS containing mutants (GSAS and double mutant 2P/GSAS) resulted in nAb titers that trended higher than the WT and 2P analogues (**Fig. 1c**). While the nAb levels from these two GSAS-containing antigens were not significantly different from one another, we believe there could be two explanations for this: first, the mutation on the furin cleavage site may alter S protein trafficking efficiency to the cell surface. Furin could be active in the trans-*golgi* network, cell surface or endosome in processing viral glycoproteins during viral maturation^49^. Thus, it is possible that blocking furin cleavage may have changed S protein trafficking from the ER to the cell surface. Although this hypothesis is unlikely, MERS and SARS, as well as other coronavirus S protein, have been reported to be missing a furin cleavage site, indicating that furin cleavage is not absolutely necessary for viral maturation^26,27,30,35^. Nonetheless, additional investigation would be needed to further understand the effect of the furin cleavage site on viral morphogenesis and SARS-CoV-2 S protein trafficking. A second possibility, which is more likely, is that we tested the mRNA vaccines at a poorly differentiating dose level (0.4 μg/dose) in mice (**Fig. 1c**). Our results in a subsequent experiment (**Fig. 2**) confirmed that the saturation point for neutralizing antibody responses in mice was between 0.2 μg and 1 μg per dose. With these considerations, we selected the double mutant formulation, MRT5500, to favor the pre-fusion form. Our dose ranging studies in NHPs confirmed the potency of MRT5500 in eliciting neutralizing antibodies. Although the sample size of our experiment (4 animals per group) was not enough to discriminate between the dose regimens, it suggested the potential of MRT5500 vaccine candidate to elicit potent neutralizing antibodies in clinic.

The long-term durability of our vaccine candidates for COVID-19 across all modalities is still under investigation. As a novel vaccine platform, mRNA can drive efficient de novo antigen expression, which is expected to activate immune responses. However, it is unknown whether the transient nature of antigen expression is sufficient in driving adequate germinal center formation which is needed for effective expansion and maturation of antigen-specific B cells. Although an mRNA vaccine for cytomegalovirus gB has demonstrated sustained antibody responses in rabbits up to 20 weeks^50^, the durability for S antigen mRNA remains an important focus for COVID-19 vaccine research. It should be noted that natural infection in COVID-19 patients, especially those of mild and asymptomatic cases, induce antibodies that decay rapidly in convalescent phase, with some drifting down to baseline within three months after diagnosis^51^. Additional preclinical studies are ongoing to further our understanding and characterization of MRT5500 and its immunological effects for applications towards COVID-19.

In summary, we have utilized mRNA technology for the rapid evaluation of vaccine candidates for COVID-19, and our results led to the selection of a double mutant candidate which has a better potential to preserve a pre-fusion conformation. The candidate MRT5500 has been shown to be immunogenic by eliciting potent neutralizing antibodies in mice and NHPs, and T_H_1-biased cellular immune responses. The candidate is positioned for further development in clinical studies as a vaccine for the prevention of COVID-19.

## Acknowledgements

We are grateful for assistance on statistical analysis by Alice Raillard and Nada Assi of Sanofi Pasteur. We also want to thank exceptional support from veterinary staff and animal research staff at New Iberia Research Center, LA and Covance, Denver, PA. The research is funded by Translate Bio and Sanofi Pasteur.

## Material and methods

### mRNA synthesis, lipid nanoparticle formulation and expression assay

Messenger RNA incorporating coding sequences containing either the wild type (WT) sequence, stabilized pre-fusion mutant (2P)^52^, furin cleavage site mutant (GSAS)^35,53^ or double mutant (2P, GSAS) of the full length SARS-CoV-2 spike glycoprotein were synthesized by *in vitro* transcription employing RNA polymerase with a plasmid DNA template encoding the desired gene using unmodified nucleotides. The resulting purified precursor mRNA was reacted further via enzymatic addition of a 5’ cap structure (Cap 1) and a 3’ poly(A) tail of approximately 200 nucleotides in length as determined by gel electrophoresis. The vaccine sequence is based on Wuhan Hu-1 strain (Genbank accession MN908947). Preparation of mRNA/lipid nanoparticle (LNP) formulations was described previously^54^. Briefly, an ethanolic solution of a mixture of lipids (ionizable lipid, phosphatidylethanolamine, cholesterol and polyethylene glycol-lipid) at a fixed lipid and mRNA ratio were combined with an aqueous buffered solution of target mRNA at an acidic pH under controlled conditions to yield a suspension of uniform LNPs. Upon ultrafiltration and diafiltration into a suitable diluent system, the resulting nanoparticle suspensions were diluted to final concentration, filtered and stored frozen at −80°C until use. Expression of S-proteins from cells transfected with synthetic mRNAs was evaluated by Western blot. Briefly, 5X10^5^ HEK293 cells were transfected using 1 μg of mRNA complexed with Lipofectamine 2000, and allowed to incubate 20 hs at 37°C. Cells were harvested after incubation period, and lysates were analyzed by Western Blot as described elsewhere^55^.

### Animal studies

Animal experiments were carried out in compliance with all pertinent US National Institutes of Health regulations and were conducted with approved animal protocols from the Institutional Animal Care and Use Committee (IACUC) at the research facilities.

The mouse studies were conducted at Covance Inc, Denver, PA. Female specific pathogen free BALB/c mice of 6-8-week-old were vaccinated in groups of 10, with 50 μL of the designated mRNA/LNP formulation into one hind leg for the prime (D0) and the contralateral hind leg for the boost (D21). Sera were collected on D-7, 14, 21, 28 and 35 from the orbital sinus or by exsanguination on D35 by the jugular vein/carotid artery. For cell-mediated response measurements, splenocytes from mice were collected on D35.

Cynomolgus macaques of Mauritian origin 2-6 years of age and weighing in a range of 2-6 kg were administered with 500 μL mRNA/LNP formulations via IM route into the deltoid of the right forelimb for the prime (D0) and the opposite forelimb for the boost (D21). Sera were collected on D-4, 14, 21, 28, 35, 42 and, PBMCs were isolated on D42. All immunizations and blood draws occurred under sedation with Ketamine HCl (10mg/kg) or Telazol (4-8mg/kg IM).

### Convalescent human sera

Convalescent serum panel (N=93) was obtained from commercial vendors (Sanguine Biobank, iSpecimen and PPD). These subjects had a PCR positive diagnosis of COVID-19 and the serum samples were collected within 3 months following diagnosis.

### Enzyme-Linked Immunosorbent Assay (ELISA)

Nunc MaxiSorb plates were coated with SARS-CoV S-GCN4 protein (custom made at GeneArt) protein at 0.5 μg/ml in PBS overnight at 4°C. Plates were washed 3 times with PBS-Tween 0.1% before blocking with 1% BSA in PBS-Tween 0.1% for 1 h at ambient temperature. Samples were plated with 1:450 initial dilution followed by 3-fold, 7-point serial dilution in blocking buffer. Plates were washed 3 times after 1-h incubation at room temperature before adding 50 μl of 1:5000 Rabbit anti-human IgG (Jackson Immuno Research) to each well. Plates were incubated at room temperature for 1hr and washed 3x. Plates were developed using Pierce 1-Step Ultra TMB-ELISA Substrate Solution for 0.1 h and stopped by TMB stop solution. Plates were read at 450 nm in SpectraMax plate reader. Antibody titers were reported as the highest dilution that is ≥0.2 Optical Density (OD) cutoff.

For mouse sera, the procedure was similar except the following differences. First, 2019-nCoV Spike protein (S1+S2) ectodomain (Sino Biological, Cat# 40589-V08B1) was used as substrate and coated at 2 μg/mL concentration in bicarbonate buffer overnight at 4°C. Second, the plates were developed using colorimetric substrate, Sure Blue TMB 1-component (SERA CARE, KPL Cat# 5120-0077) and stopped by Stop solution (SERA CARE Sure Blue, KPL Cat# 5120-0024). The endpoint antibody titer for each sample was determined as the highest dilution which gave OD value 3x higher than the background.

### Pseudovirus Neutralization Assay

Serum samples were diluted 1:4 in media (FluoroBrite phenol red free DMEM +10% FBS +10mM HEPES +1% PS + 1% Glutamax) and heat inactivated at 56°C for 0.5 h. A further, 2-fold serial dilution of the heat inactivated serum were prepared and mixed with the reporter virus particle (RVP) -GFP (Integral Molecular) diluted to contain 300 infectious particles per well and incubated for 1 h at 37°C. 96-well plates of 50% confluent 293T-hsACE2 clonal cells in 75 μL volume were inoculated with 50 μL of the serum/virus mixtures and incubated at 37°C for 72h. At the end of the incubation, plates were scanned on a high-content imager and individual GFP expressing cells were counted. The inhibitory dilution titer (ID_50_) was reported as the reciprocal of the dilution that reduced the number of virus plaques in the test by 50%. ID_50_ for each test sample was interpolated by calculating the slope and intercept using the last dilution with a plaque number below the 50% neutralization point and the first dilution with a plaque number above the 50% neutralization point. ID_50_ Titer = (50% neutralization point - intercept)/slope).

### Microneutralization assay

Serial two-fold dilutions of heat inactivated serum samples were incubated with a challenge dose targeting 100 50% tissue culture infectious dose (TCID_50_) of SARS-CoV-2 (strain USA-WA1/2020 [BEI Resources; catalog# NR-52281]) at 37°C with 5% CO_2_ for 1 hour (h). The serum-virus mixtures were inoculated into wells of a 96-well microplate with preformed Vero E6 (ATCC® CRL-1586^TM^) cell monolayers and adsorbed at 37°C with 5% CO_2_ for 0.5 h. Additional assay media was added to all wells without removing the existing inoculum and incubated at 37°C with 5% CO_2_ for 2 days. After washing and fixation of the Vero E6 cell monolayers, SARS-CoV-2 antigen production in cells was detected by successive incubations with an anti-SARS-CoV nucleoprotein mouse monoclonal antibody (Sino Biological catalog# 40143-MM05), HRP IgG conjugate (Jackson ImmunoResearch Laboratories, catalog #115-035-062), and a chromogenic substrate. The resulting optical density (OD) was measured using a microplate reader. The reduction in SARS-CoV-2 infectivity, as compared to that in the virus control wells, constitutes a positive neutralization reaction indicating the presence of neutralizing antibodies in the serum sample. The 50% neutralization titer (MN_50_) was defined as the reciprocal of the serum dilution for which the virus infectivity was reduced by 50% relative to the virus control on each plate. The MN_50_ for each sample was interpolated by calculating the slope and intercept using the last dilution with an OD below the 50% neutralization point and the first dilution with an OD above the 50% neutralization point; MN_50_ Titer = (OD of 50% neutralization point - intercept)/slope.

### Cytokine ELISPOT analysis

For testing cytokine responses in mice CTL ELISPOT kits (Mouse IFN-γ/IL-5 Double-Color enzymatic ELISPOT, Immunospot) were used according to the manufacture’s protocols. Briefly, freshly isolated splenocytes were resuspended in CTL-Test Media and incubated overnight at 300,000 cells per well with commercially available SARS-CoV-2 S peptide pools. PepMix™ SARS-CoV-2 (Spike Glycoprotein, Cat# PM-WCPV-S-1, JPT, Germany) peptide pool 1 and pool 2 were used at the final concentration of 2 μg/ml per well. Concanavalin A (CovA, Sigma C5275) at concentration of 1 μg/ml was used for a positive control stimulation. After overnight incubation, the plates were washed and developed per manufacturer instructions. Spots were scanned and analyzed by the CTL technical team. The number of cytokines producing cells per million cells was reported.

For testing cytokine responses in NHPs Monkey IFNɣ ELISPOT (CTL, cat# 3421M-4APW) and IL-13 ELISPOT kits (CTL, cat# 3470M-4APW) were used. Previously frozen PBMCs were washed, resuspended in culture medium provided by the kit and enumerated. PepMix™ SARS-CoV-2 peptide pools as well as CovA were used for stimulation as described above. PBMC were plated at 300,000 cells per well and stimulated overnight. After overnight incubation the plates were washed and developed per manufacturer instructions. The plates were dried overnight, scanned, and spots were counted using a CTL analyzer (Immunospot S6 Universal Analyzer, CTL). The data were reported as spot forming cells (SFC) per million PBMCs.

### Statistical analysis

Data were log-10 transformed prior to statistical analysis. All statistical tests were two-sided, and the nominal level of statistical significance was set to α=5%. All analyses were performed on SEG SAS v9.4®. Statistical comparisons among different groups (different dose levels or constructs in a particular study) or between D35 and pre-bleed were conducted using mixed effect model for repeated measures, the model included group, day and their interactions, where day was specified as repeated measures.

When assessing pairwise correlations among IgG, MN, PsVNa in NHP study, we proposed a two-stage approach to separate the intra-and inter-variabilities for the repeated measures. Stage 1: we calculated the correlation coefficient for each individual subject based on observations over time per subject; Stage 2: we then estimated the mean and 95% CI of group correlation coefficient based on individual coefficient estimates. The analysis was based on log 10 transformed data.

Statistical comparisons among different groups (i.e. different dose levels) and the convalescent sera on D35 were conducted using either analysis of variance (ANOVA) or Wilcoxon Rank Sum Test.

