## Supplemental materials for "Immunogenicity of novel mRNA COVID-19 vaccine MRT5500 in mice and non-human primates"

^1^Sanofi Pasteur, 38 Sidney Street, Cambridge, MA 02139, ^2^Sanofi Pasteur, Discovery Dr., Swiftwater, PA 18370, ^3^Translate Bio, 29 Hartwell Ave, Lexington, MA 02421, ^4^Sanofi Pasteur, [1541 AV Marcel Mérieux, 69280 Marcy l'Etoile](https://eur01.safelinks.protection.outlook.com/?url=https:%2F%2Fwww.bing.com%2Flocal%3Flid%3DYN2000x670688274%26id%3DYN2000x670688274%26q%3DSanofi%2BPasteur%2B(S.A)%26name%3DSanofi%2BPasteur%2B(S.A)%26cp%3D45.77716827392578~4.706181049346924%26ppois%3D45.77716827392578_4.706181049346924_Sanofi%2BPasteur%2B(S.A)&data=02%7C01%7Ckirill.Kalnin%40sanofi.com%7C92f0fa45cd2440a4cd9f08d8694603dd%7Caca3c8d6aa714e1aa10e03572fc58c0b%7C1%7C0%7C637375096712767399&sdata=bEhPKQGY7aGMddSkCG5HpEuPVX1vuqms0vfB%2FpYgTwo%3D&reserved=0), France, ^5^Yoh Services LLC, 38 Sidney Street, Cambridge, MA 02139

**Summary**

An effective vaccine to address the global pandemic of coronavirus disease 2019 (COVID-19) is an urgent public health priority^1^. Novel synthetic mRNA and vector-based vaccine technologies offer an expeditious development path alternative to traditional vaccine approaches. Here we describe the efforts to utilize an mRNA platform for rational design and evaluations of mRNA vaccine candidates based on Spike (S) glycoprotein of Severe Acute Respiratory Syndrome Coronavirus 2 (SARS-CoV-2), the virus causing COVID-19. Several mRNA constructs expressing various structural conformations of S-protein, including wild type (WT), a pre-fusion stabilized mutant (2P), a furin cleavage-site mutant (GSAS) and a double mutant form (2P/GSAS), were tested in a preclinical animal model for their capacity to elicit neutralizing antibodies (nAbs). The lead 2P/GSAS candidate was further assessed in dose-ranging studies in mice and *Cynomolgus* macaques. The selected 2P/GSAS vaccine formulation, now designated MRT5500, elicited potent nAbs as measured in two types of neutralization assays. In addition, MRT5500 elicited T_H_1-biased responses in both mouse and non-human primate species, a result that helps to address a hypothetical concern regarding potential vaccine-associated enhanced respiratory diseases associated with T_H_2-biased responses. These data position MRT5500 as a viable vaccine candidate for clinical development against COVID-19.

**Key words:** COVID-19, SARS-CoV-2, vaccine, mRNA, LNP, BALB/c mice, cynomolgus macaques, immunogenicity, neutralization potency, neutralization, microneutralization, ELISA

**Supplemental materials**

**Supplemental Figure 1.** MRT5500 elicited strong binding anti-spike response in NHPs (see Materials and Methods)

**Supplemental Figure 2**: MRT5500 induces T_H_1-biased T-cell responses in mice. D35 (A) IFNɣ and (B) IL-5 ELISPOT data for the 5 and 10 µg dose groups following the overnight re-stimulation with S-protein peptide poolles. Testing was performed on splenocyte pools within the group. The frequencies of immune splenocytes secreting IFNɣ or IL-5 were calculated as spots forming cells (SFC) per million cells.

**Supplemental Figure 3.** Strong correlations between individual NHP ELISA, PsV and MN time-point titers (see **Suppl Table 1**). Top panel A: 4 subjects in 15 µg dose; Middle panel B: 4 subjects in 45 µg dose; Bottom panel C: 4 subjects in 135 µg dose.

**Supplemental Figure 4.** PsV titers in mice for the 1 μg, 5 μg and 10 μg dose levels of MRT5500 were significantly different from the Human Convalescent sera PsV titers. The horizontal line in the boxplot indicates the median, the box is the interquartile range and the whiskers is the range.

**Supplemental Figure 5.** MRT5500 MN (a) and PsV (b) titers in NHPs were significantly different from MN and PsV titers of Human Convalescent Sera. The horizontal line in the boxplot indicates the median, the box is the interquartile range and the whiskers is the range.

**Supplemental Table 1.** Spearman Correlation Coefficients (SCC) between ELISA (IgG), Pseudoviral (PsV) and Microneutralization (MN) titers. SCC were conducted per individual animals (Suppl. Fig.4) and Means (95% CI) were calculated per dose (N=4) or all NHPs (N=12)

**Supplemental Table 2.** Pairwise dose comparison in PsV neutralization titers on D35 in mice. There were no statistically significant differences in PsV titers among the 1 μg, 5 μg and 10 μg dose levels, while at the lowest dose level (0.2 μg) PsV titers were significantly different from those obtained with the higher dose levels.

**Supplemental Figures**

**Supplemental Figure 1**

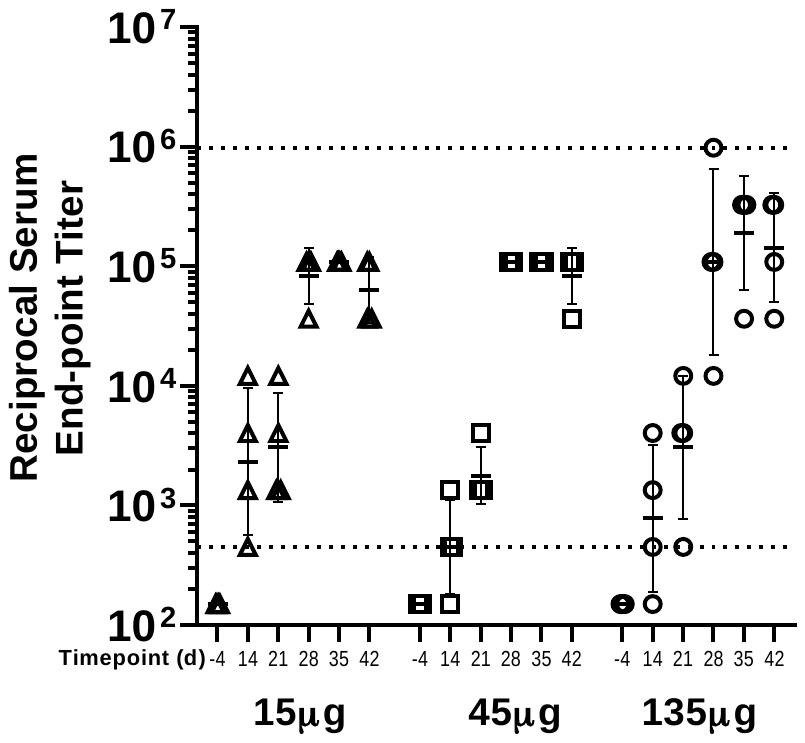

**Supplemental Figure 2**
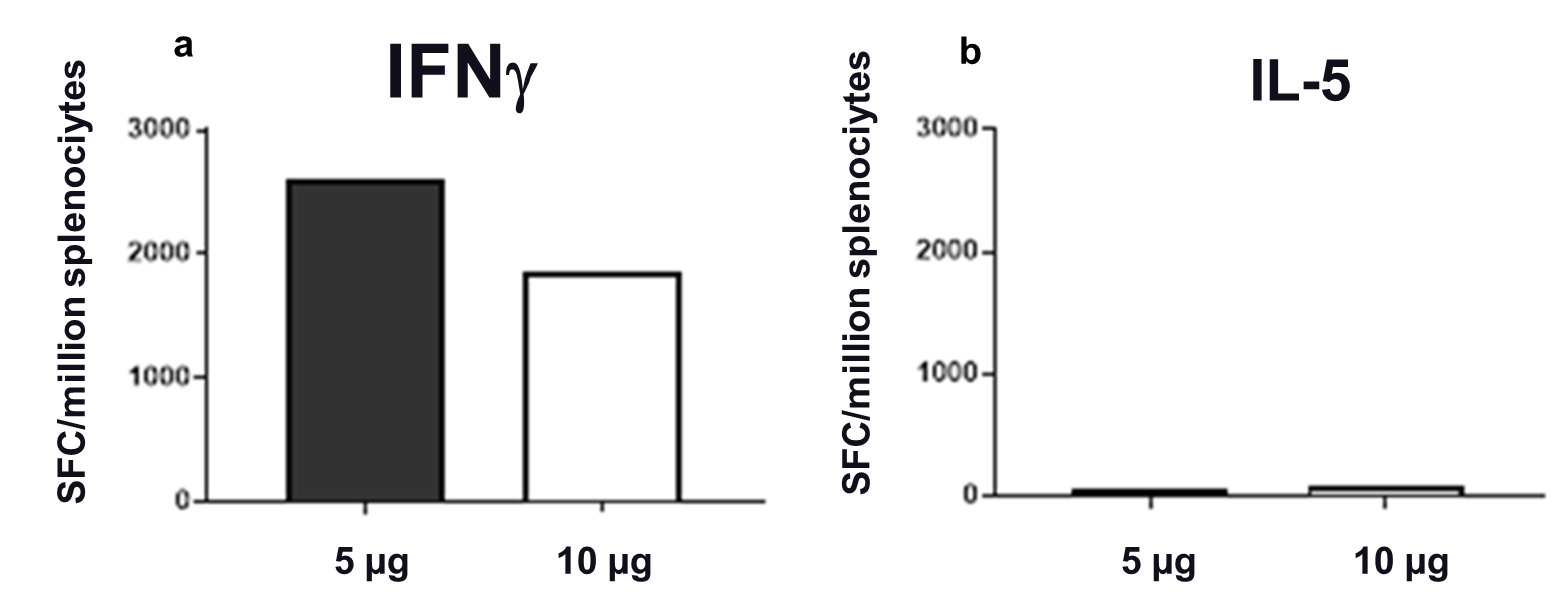

**Supplemental Figure 3**
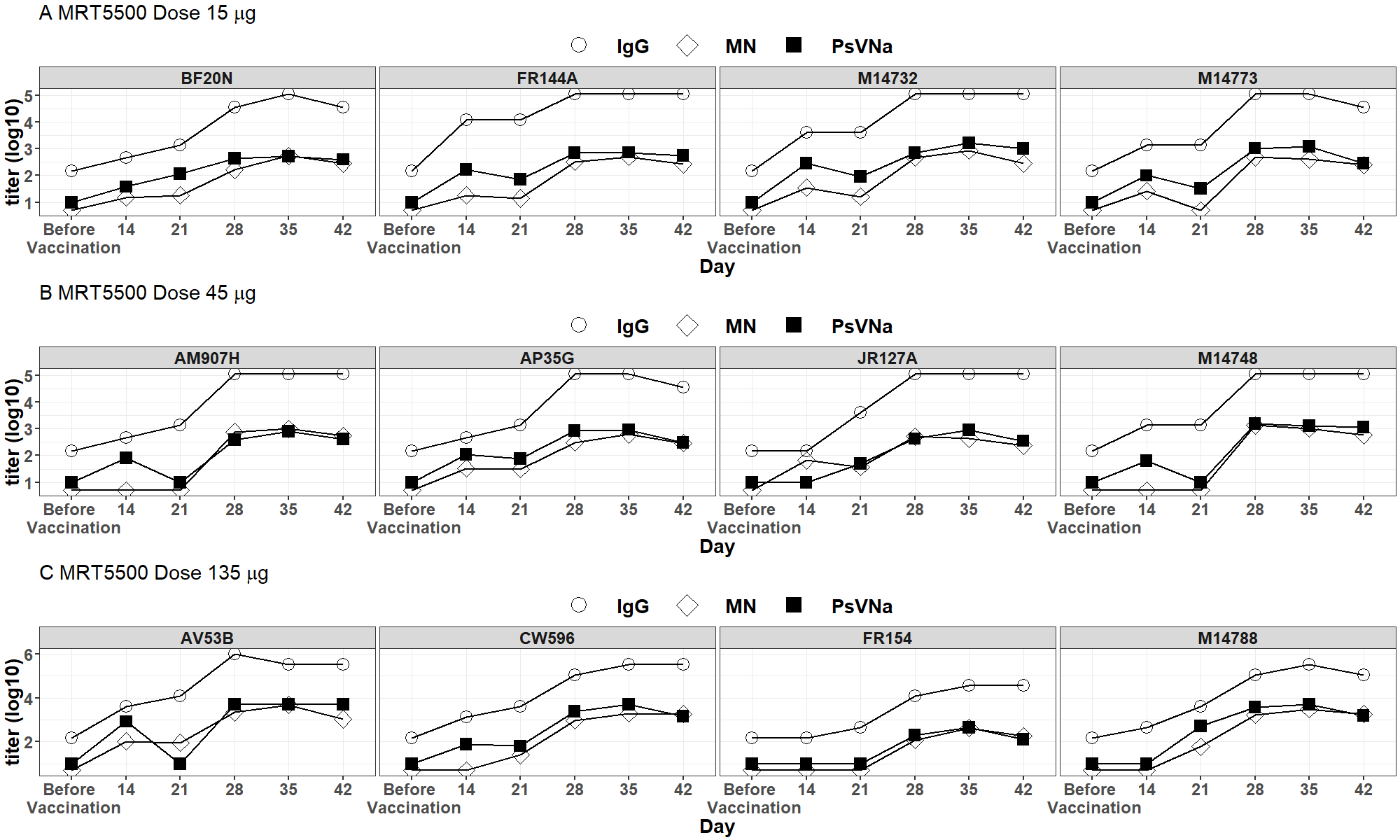

**Supplemental Figure 4**

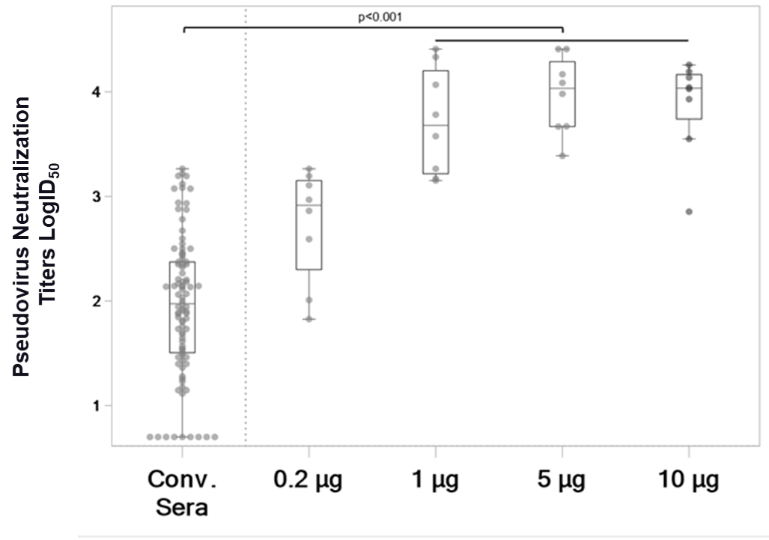

**Supplemental Figure 5**
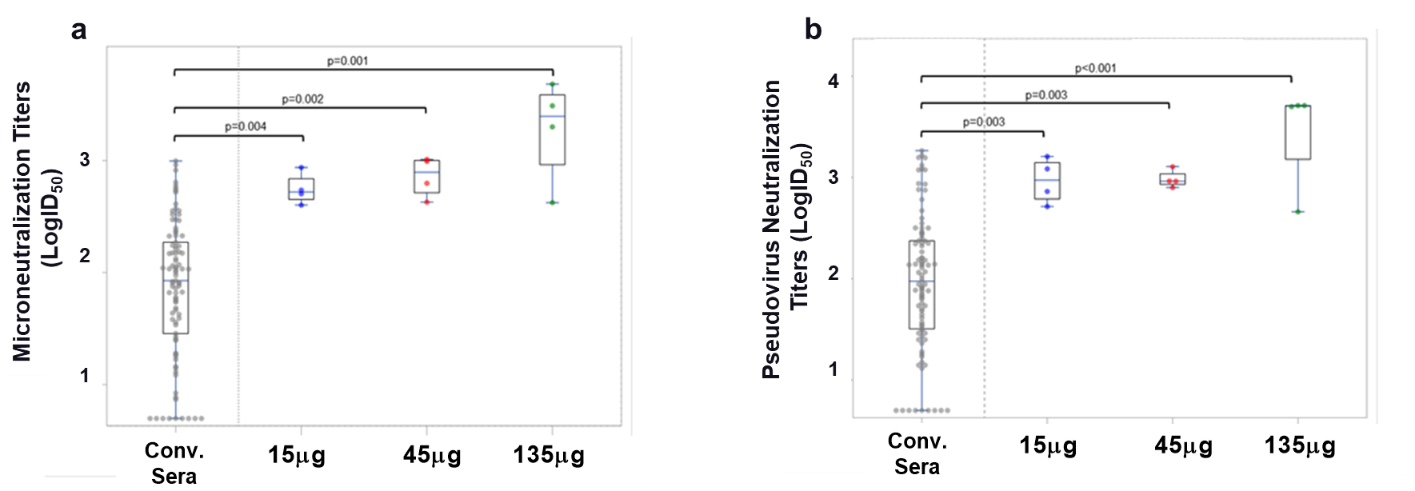

**Supplemental Table 1.**

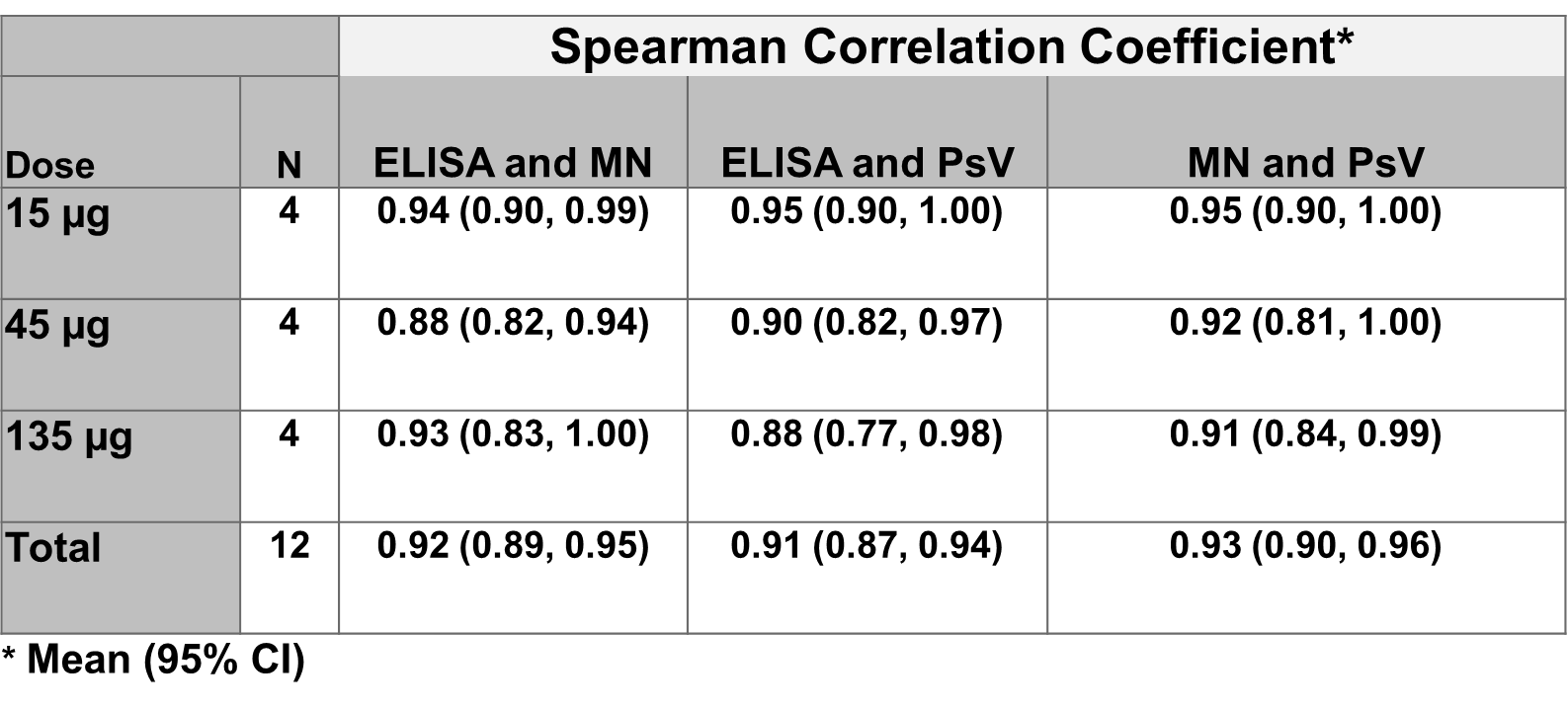

**Supplemental Table 2.**

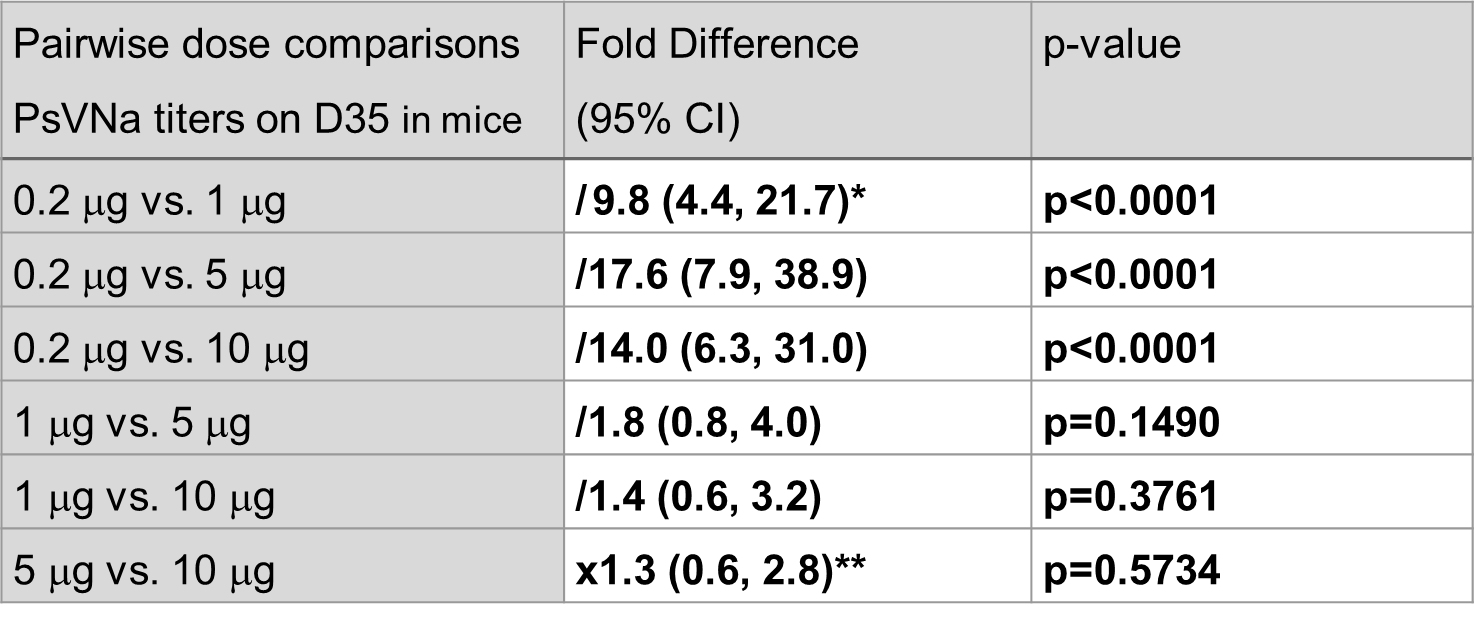

Comparison: Group1 versus Group2
* /X.X: Group1 is X.X-fold lower than group2

** xX.X: Group1 is X.X-fold higher than group21	Graham, B. S. Rapid COVID-19 vaccine development. *Science* **368**, 945-946, doi:10.1126/science.abb8923 (2020).
